# Predicting Primary Site of Secondary Liver Cancer with a Neural Estimator of Metastatic Origin (NEMO)

**DOI:** 10.1101/689828

**Authors:** Geoffrey F. Schau, Erik A. Burlingame, Guillaume Thibault, Tauangtham Anekpuritanang, Ying Wang, Joe W. Gray, Christopher Corless, Young Hwan Chang

## Abstract

Pathologists rely on clinical information, tissue morphology, and sophisticated molecular diagnostics to accurately infer the metastatic origin of secondary liver cancer. In this paper, we introduce a deep learning approach to identify spatially localized regions of cancerous tumor within hematoxylin and eosin stained tissue sections of liver cancer and to generate predictions of the cancer’s metastatic origin. Our approach achieves an accuracy of 90.2% when classifying metastatic origin of whole slide images into three distinct classes, which compares favorably to an established clinical benchmark by three board-certified pathologists whose accuracies ranged from 90.2% to 94.1% on the same prediction task. This approach illustrates the potential impact of deep learning systems to leverage morphological and structural features of H&E stained tissue sections to guide pathological and clinical determination of the metastatic origin of secondary liver cancers.

## 1 Introduction

Metastatic liver cancer accounts for 25% of all metastases to solid organs, yet because liver metastases can arise from almost anywhere in the body, accurately determining the origin of metastatic liver cancer is of paramount importance for guiding effective treatment.^1, 2^ In clinical practice, pathologists commonly rely on clinical information, tissue examination, and molecular assays to determine the metastatic origin of a patient’s secondary liver tumor. Although clinically effective, this approach requires significant expertise, experience, and time to perform properly.

Deep learning methods have rapidly accelerated the automation of key processes in identifying and quantifying clinically meaningful features in biomedical images and continues to drive modern advancements in digital pathology.^3, 4^ Furthermore, deep learning systems have been applied to settings where their performance matches and even exceeds the ability of clinical human practitioners in tasks related to image analysis, including in clinical instances that rely on inspection of hematoxylin and eosin (H&E) stained tissue.^5–9^ The emerging power and success of many deep learning approaches applied to image content analysis stem from their ability to learn and leverage meaningful features from large data data collections that cannot be explicitly mathematically modeled.^6, 10–12^ For example, these approaches can provide robust and reproducible solutions for automated detection and analysis of tumor lesions within whole slide images containing both normal and cancerous tumor tissue segments.^13–15^

Our key contribution in this paper is a deep learning approach to identify metastatic tissue within whole slide section and classify these tumors by their metastatic origin. We evaluate model performance with respect to a clinical benchmark established by three board-certified pathologists charged with the same classification task as our model in which each pathologist was tasked to infer the metastatic origin of liver cancer directly from H&E stained tissue sections without the use of molecular immunohistochemistry assays or clinical data. Through this work, we demonstrate feasibility of deep learning systems to automatically characterize the biological origin of metastatic cancers by their morphological features presented in H&E tissue sections.

## 2 Method

Our approach is composed of two deep neural networks that operate in series. The first stage model is trained to filter tiles containing normal or stromal tissue from whole slide images (WSIs), as these tiles are not expected to have predictive value in estimating the metastatic origin of cancerous tissue. A second stage model is then trained to predict a single label of metastatic origin for each tile in the dataset and aggregate tile predictions within independent WSIs to generate a single whole-slide prediction of metastatic origin. A diagram illustrating the basic workflow of our approach is shown in Fig. 1.

**Fig 1.**
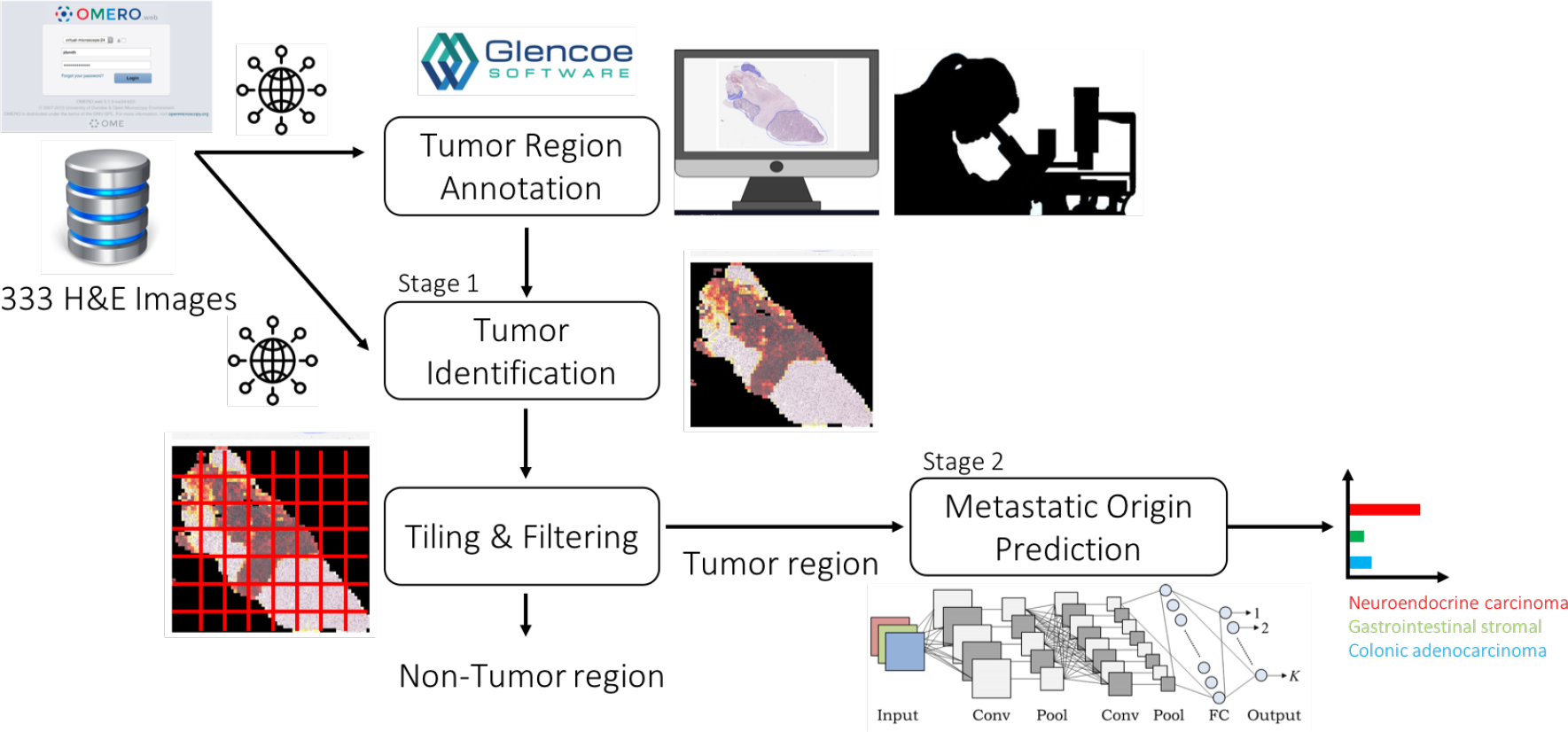
Deep learning based approach to leverage pathological annotation of tumor region to isolate and localize tumor tissue from a WSI and generate predictions of metastatic origin.

Our approach first leverages pathologist annotation of tumor region to train a binary classifier designed to predict whether a given tile of H&E image is either tumor or non-tumor tissue. A second stage classification model is then trained on the tumor portions of images to correctly predict their metastatic origin with respect to clinically-determined labels. In all cases, model output is reassembled into probabilistic heatmaps over the WSI, enabling a rapid assessment of spatial characteristics driving predictive reasoning. Both first and second stage models utilize the Inception v4 deep learning architecture,^16^ which is optimized to capture morphological and architectural features on varying scales with high efficiency and has been shown to achieve human-level prediction capability on the ImageNet dataset. Models were developed in Keras with Tensorflow backend^17^ and trained undergoing cyclic learning rates^18^ using the Adam optimizer^19^ on NVIDIA V100 GPUs made available through the Exacloud HPC resource at Oregon Health & Science University.

Raw H&E images were acquired from the OHSU Knight BioLibrary, uploaded to a secure instance of an OMERO server,^20^ programmatically accessed through the OpenSlide python API,^21^ normalized with established methods to overcome known inconsistencies in the H&E staining process,^22^ and tiled into non-overlapping patches of 299 *×* 299 pixels necessary to accommodate the Inception v4 architecture. Tiles whose mean three-channel, 8-bit intensities were greater than 240 were filtered out as white non-informative background. Pathological annotations of tumor regions within 28 whole slide H&E images from each of the fourteen metastatic subtypes were collected using PathViewer (https://glencoesoftware.com/products/pathviewer/), an interactive utility for the collection and storage of pathological annotation. Tumor region annotations defined the target label for each tile in the WSI dataset as either belonging to tumor or non-tumor tissue. We randomly assigned 30% of whole slide images to a held-out test set used for model validation. The code used to generate the results and figures is available in a dedicated Github repository.

## 3 Results

### 3.1 Quantitative Localization of Liver Cancer in Whole Slide Images

The first-stage model is a tumor tile binary classifier that generates a prediction between 0 and 1 for each tile in the dataset in which a 1 corresponded to perfect confidence that a tile was of tumor tissue and in which a 0 corresponded to perfect confidence that the tile was of normal or stomal tissue. This model achieved an AUC of 0.74 under the receiver operator characteristics curve which was sufficient to establish good correlation (*R*^2^ = 0.96) between clinical estimation and our model’s estimates of tumor purity in whole slide H&E images as shown in Fig. 2. Further, visual comparisons between the pathological tumor annotation and our model’s predictions illustrate spatial concordance between the drawn tumor-bounding mask and our model’s predictions. Once trained, the tumor-region identifying model was deployed on the entire remaining dataset to include only tiles containing cancerous tissue.

**Fig 2.**
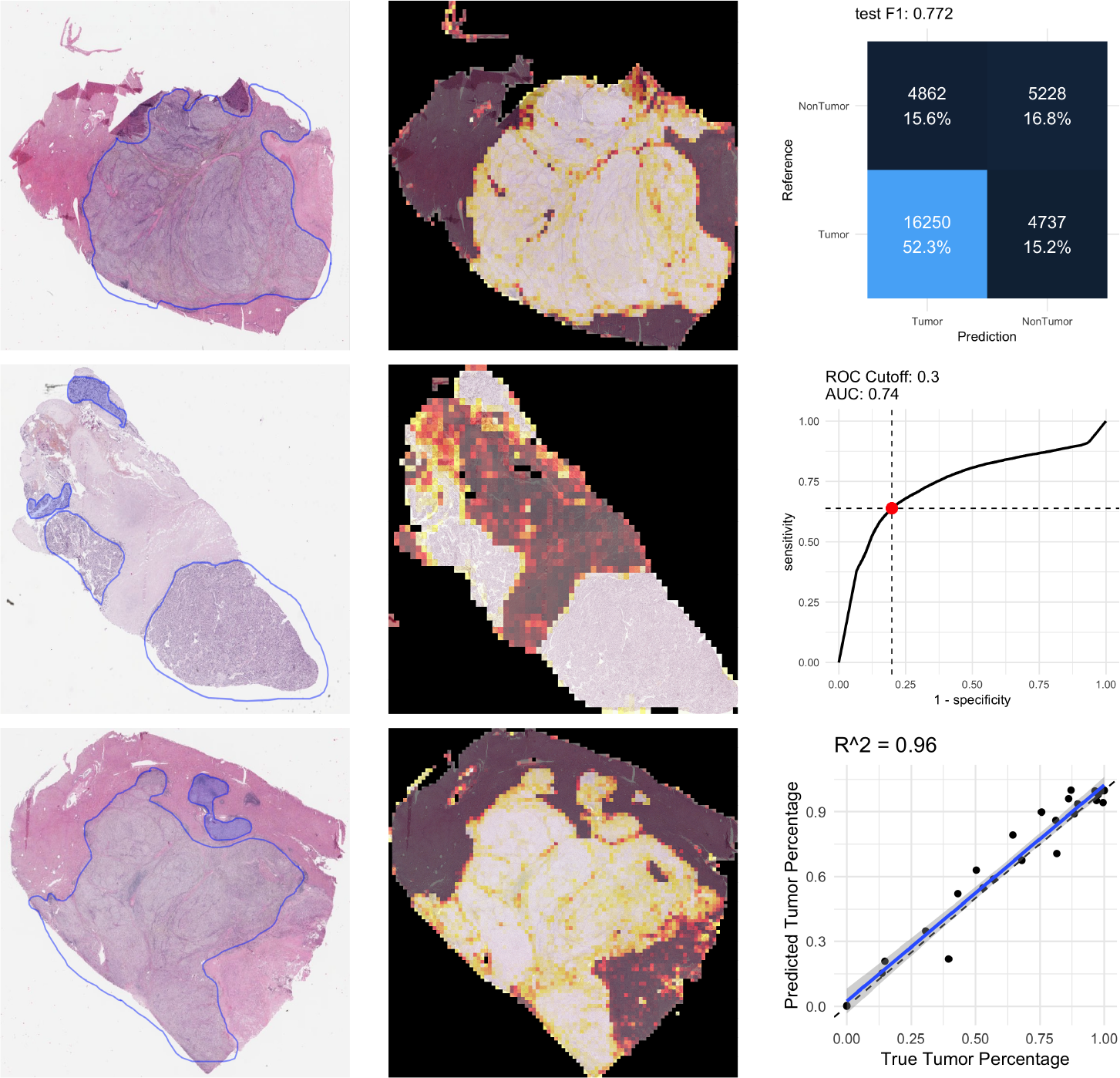
(left column) Three examples from the held-out testing set with pathological annotation of tumor regions outlined in blue. (center column) Corresponding model predictions estimating regions of whole slide images that contain tumor tissue show concordance between the pathological annotation of tumor region with the outcome of our model. In these illustrations, a brighter color intensity corresponds to higher probability that the underlying tile was labeled as being of tumor by the trained model. (right column, top) Confusion matrix from the held-out testing set for a tumor/non-tumor predictive model illustrating F1 score of 0.772 in the classification task. (right column, center) Receiver Operating Characteristics (ROC) curve illustrating area under the curve of 0.74. (right column, bottom) Comparison between the true tumor purity in the sample inferred from the pathological annotation (x-axis) versus the inferred tumor purity from the model’s output (y-axis) with strong correlation (*R*^2^ = 0.96)

### 3.2 Quantitative Whole Slide Image Classification of Metastatic Origin

Each image in the dataset is annotated with clinically-determined metastatic origin labels informed by clinical information, pathological inspection of tissue sample, and IHC profiling. These clinical annotations were summarized into 14 distinct subgroups by a clinical practitioner, which are shown in Fig. 3. Although exploratory data analysis considered all fourteen classes, due to class imbalances in this dataset, this work is predicated on the three primarily represented metastatic origins in the dataset (colonic adenocarcinomas, neuroendocrine carcinomas, and gastrointestinal stromal tumors) which collectively represent over 80% of the relevant data.

**Fig 3.**
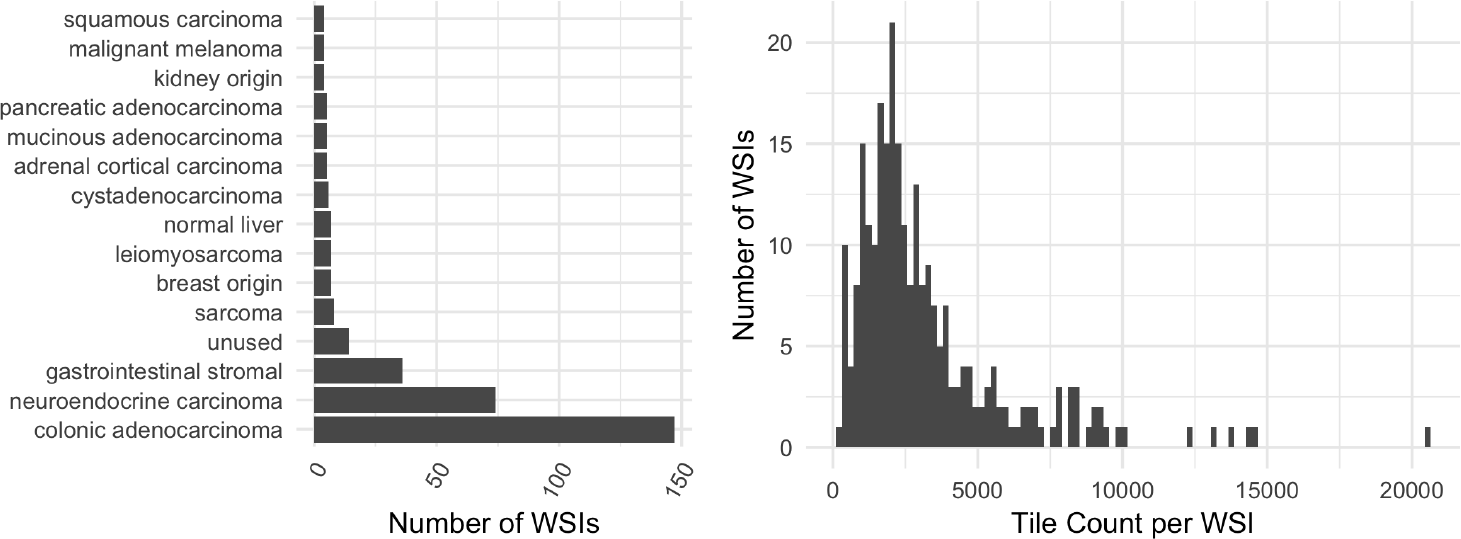
(left) Summary of the acquired dataset, composed of 333 WSIs each containing metastatic tissue originating from one of 14 sites. (right) Distribution of non-overlapping tile counts in each WSI with mean count 3302 tiles per WSI.

#### 3.2.1 Whole Slide Image Classification

After the first stage identifies regions of the H&E images that are tumor, the second stage model learns to classify those tiles according to their metastatic origin. A second Inception v4 deep neural network was designed to generate a three-class prediction for each tile in the training set as belonging to either a colonic adenocarcinoma, gastrointestinal stromal, or neuroendocrine carcinoma. Whole slide image predictions predictions aggregated across all corresponding tumor tiles achieved an F1 score of 0.875 on the held-out testing set of WSIs, having failed to correctly classify 5 out of the 51 held-out testing samples. Class-specific statistics shown in Table 1 quantify classification performance metrics for the metastatic origin prediction model. Confusion matrices of both WSI and per-tile predictions are shown in Fig. 4.

**Table 1.** Class-specific statistics of both the tumor identification and three-way origin classification task

|  | Sensitivity | Specificity | Pos Pred Value | Neg Pred Value | Precision | Recall | F1 |
| --- | --- | --- | --- | --- | --- | --- | --- |
| Tumor Identification | 0.77 | 0.52 | 0.77 | 0.53 | 0.77 | 0.77 | 0.77 |
| Colonic Adenocarcinoma | 1.00 | 0.83 | 0.88 | 1.00 | 0.88 | 1.00 | 0.93 |
| Neuroendocrine Carcinoma | 0.79 | 1.00 | 1.00 | 0.92 | 1.00 | 0.79 | 0.88 |
| Gastrointestinal Stromal | 0.78 | 0.98 | 0.87 | 0.95 | 0.88 | 0.78 | 0.82 |

**Fig 4.**
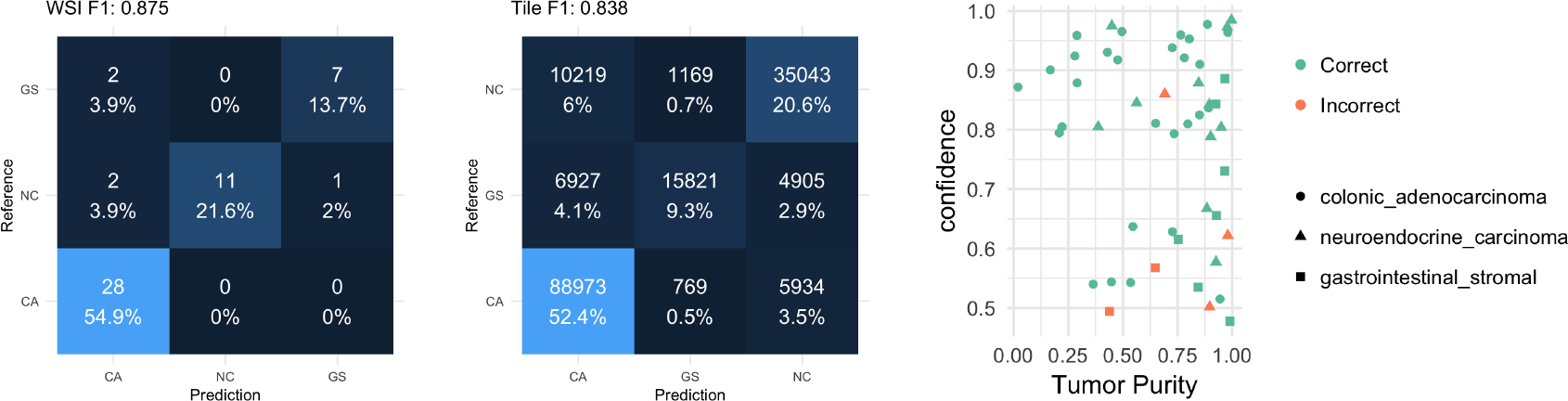
(left) Confusion matrix of WSI prediction on a held-out test set. (center) Confusion matrix of tile-based predictions. (right) failure cases with respect to the inferred tumor purity in the sample on the x-axis (fraction of tiles predicted to be tumor) and the model’s output confidence in its prediction on the y-axis.

Several technical factors were associated with incorrect predictions, including slide blurring, tissue folding, and low tumor purity. Our model’s confidence was lower for samples that it incorrectly classified, as shown in Fig. 4, though one sample was incorrectly classified with 86% confidence which was driven by misclassified stromal tissue present in teh H&E slide. Individual tiles associated with highly confident predictions for each class are shown in Fig. 5. Pathological inspection of these tiles suggests that tiles associated with highly confident class predictions presentpathological features that guide diagnoses, as the first row contains tiles presenting features associated with primarily spindle-type gastrointestinal stromal tumors and the third row presenting typical well-differentiated neuroendocrine carcinomas. The first two images in the second row represent dirty necrotic tissue which, among the three diseases under consideration, tends to be associated with colonic adenocarcinomas. However, this type of feature is not explicitly associated with cancer, and so should be interpreted with caution. Importantly, this approach obviates the need for pathological region annotation beyond what was required to train the first stage model.

**Fig 5.**
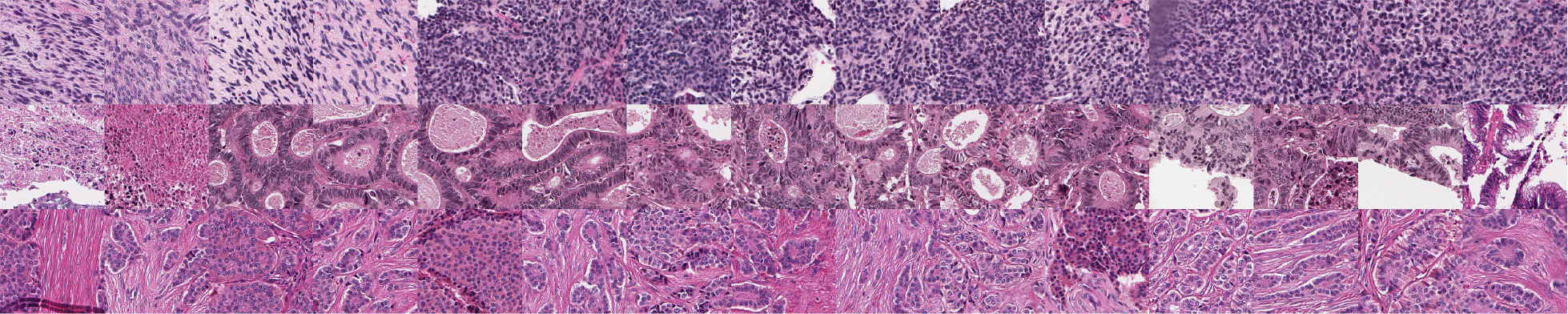
Example tiles correctly classified by the model with high confidence in which each row is a distinct class (gastrointestinal stromal, colonic adenocarcinoma, and neuroendocrine carcinoma in rows 1, 2, and 3, respectively).

### 3.3 Clinical Benchmark Comparison Study

A study was developed to benchmark our approach to clinical practitioners. This study recruited three board-certified pathologists to independently classify each of the 51 whole slide image samples in the held-out test set according to their metastatic origin. Each participant independently incorrectly classified 3, 4, and 5 samples each, while our neural network model missed 5 samples from the held-out test set. Table 2 summarizes the eleven samples that were missed by either the model or by at least one pathologist and their respective predictions. Interestingly, only two of the mis-classified samples by the model were correctly classified by all three pathologists. Figure 6 illustrates a selected sample classified correctly by the model and all three pathologists, a sample missed by the model that the pathologists all got correct, a sample missed by both the model and at least one pathologist, and a sample for which the model was correct but at least one pathologist made an incorrect classification. All examples illustrate the raw H&E image and three heatmaps generated by the model for each of the three-way predictions in which a brighter color corresponds to a higher confidence in the model’s prediction for each class. Importantly, predictions are only available for tiles that the first-stage of our model classified as tumor tissue, as non-tumor tiles were filtered out of the metastatic origin prediction task. Although the failure cases are diverse, probabilistic overlays of metastatic origin prediction may facilitate faster and more efficient examination of these tissue sections in clinical decision making processes.

**Table 2.** Slides mis-classified by either the model or at least one pathologist (GS: Gastrointestinal stromal; CA: Colonic adenocarcinoma; NC: neuroendocrine carcinoma)

| Slide Alias | Ground Truth | Model | Path1 | Path2 | Path3 |
| --- | --- | --- | --- | --- | --- |
| 101 | CA | CA | CA | CA | NC |
| 102 | CA | CA | CA | CA | NC |
| 103 | CA | CA | CA | NC | GS |
| 104 | CA | CA | NC | NC | CA |
| 105 | GS | CA | GS | GS | GS |
| 106 | GS | GS | GS | GS | NC |
| 107 | GS | CA | GS | GS | GS |
| 108 | NC | GS | GS | NC | GS |
| 109 | NC | CA | NC | NC | NC |
| 110 | NC | NC | NC | CA | NC |
| 111 | NC | CA | CA | CA | NC |

**Fig 6.**
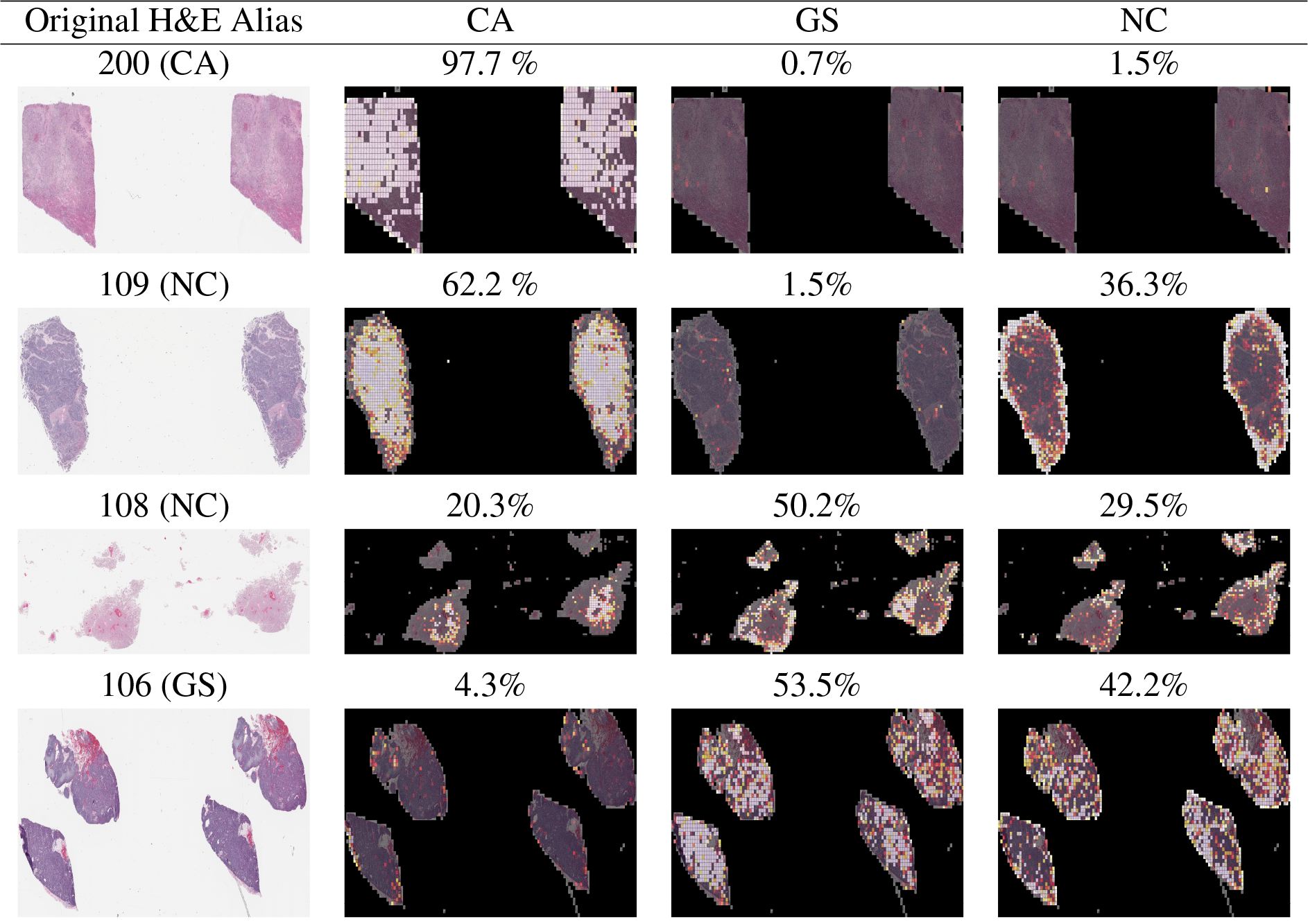
Example mis-classified H&E slides with associated annotations from the second stage model illustrating localized predictions of metastatic origin in which brighter colors are associated with more confident class-specific predictions. First row: sample correctly predicted by the model and all three pathologists. Second row: sample missed by the model that all three pathologists got correct. Third row: Example missed by both the model and at least one pathologist. Fourth row: example missed by at least one pathologist that the model got correct. (GS: Gastrointestinal stromal; CA: Colonic Adenocarcinoma; NC: Neuroendocrine Carcinoma)

#### 3.3.1 Complementary Study: Unsupervised Discovery of Origin-Related Features

As part of our exploratory data analysis, we sought to evaluate whether an unsupervised learning model can learn to identify spatial features of H&E stained tissue tiles that correspond to their metastatic origin. To guide this approach, we employed variational autoencoders (VAE) to learn latent feature representations of metastatic cancers of diverse origins.

VAEs are composed of complementary encoder and decoder networks in which the encoder network θ learns to compress an input data sample into a learned latent representation while the decoder network *ϕ* learns to reconstruct the original input from the latent code. VAEs learn unsupervised features by minimizing a composite loss function composed of two terms during training. Equation 1 illustrates that VAEs learn to minimize the difference between input sample *x* and its reconstruction as well as minimize the Kullback-Leibler (KL) divergence of the latent variable *z* distribution with respect to a known prior *p*(*z*), which in this case is the standard normal distribution. By penalizing divergence between an learned feature distribution and an expected prior, VAEs learn optimal encodings that conform to expected distributions across the dataset and enable a probabilistic interpretation of the embedded feature space.

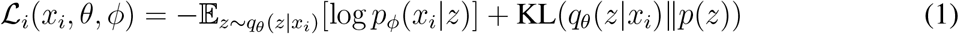

A single VAE model was trained to learn latent feature sets of individual tiles composed of four thousand tiles sampled randomly from the fourteen classes presented in the dataset. The learned features space of individual tiles are projected into two-dimensional space with the t-SNE algorithm^23^ and visualized by their respective metastatic origin in Fig. 7. This unsupervised approach suggests separability of image tiles based on features associated with metastatic origin, suggesting feasibility of a supervised classification model to correctly classify the origin of whole slide H&E images of liver metastases. Further, the learned feature space map generates a compelling arrangement of individual tiles grouped by their local feature space similarity. A complementary figure that illustrates an arrangement of tile images onto their respective coordinate projection is available online through FigShare at 10.6084/m9.figshare.8340581.

**Fig 7.**
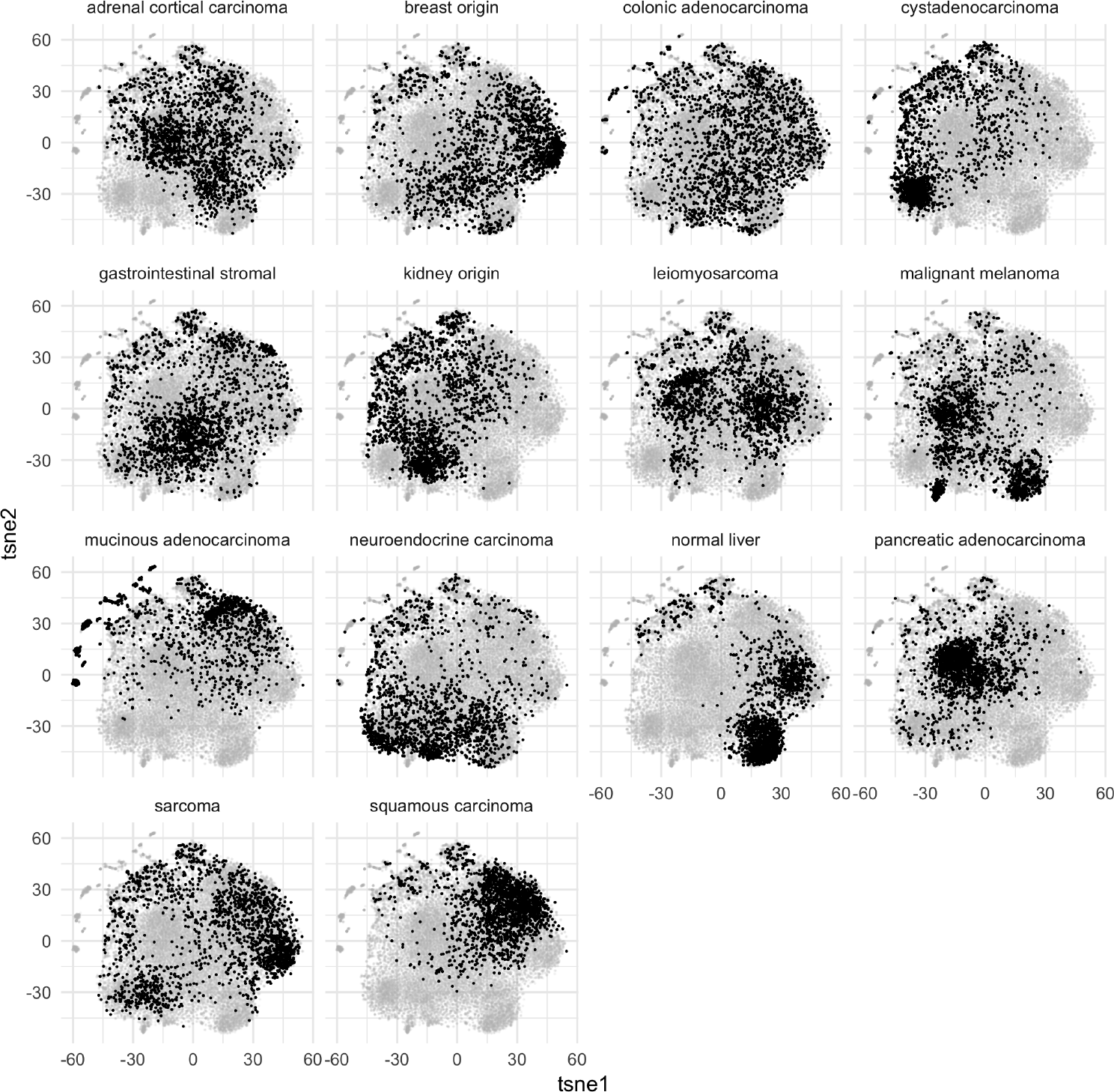
Learned unsupervised features of H&E tile images from a VAE model projected into two dimensional space and colored by the tiles’ metastatic origin illustrate local feature similarity with respect to metastatic origin. Each facet image contains the t-SNE projection of the latent space encoding of each of the tiles in the dataset. That tiles of similar metastatic origin tend to cluster together suggests that morphological features contained within the tiles are informative for estimating where the metastases originated within the body. A higher-resolution figure illustrating input tiles on their respective coordinates is hosted on FigShare (10.6084/m9.figshare.8340581).

## 4 Discussion

This work presents a deep learning based approach designed to predict the origin of metastatic liver cancer using a two-stage serial model composed of a first model trained to identify tumor from non-tumor within H&E sections of metastatic liver tissue based on pathologists annotation and a second stage model that learns to predict the metastatic origin of individual patches of tumor tissue and aggregates those results into predictions over WSIs. We illustrate through a clinical benchmark comparison that our approach is within performance criteria of board-certified pathologists, suggesting that these types of systems may be capable of generating rapid, first-pass assessments of metastatic origin in the absence of detailed clinical information or comprehensive molecular profiling assay. We believe this type of data-driven visualization augmentation provides an additional layer of information that may facilitate the speed and ease of generating final decisions by clinical care providers.

Although these results illustrate feasibility of our approach, several significant limitations remain. Principally, this analysis was data-limited to only three most-prevalent sources of metastatic origin when in practice metastases can and do originate from a broad variety of biological sources. A future direction will seek to leverage H&E stained tissue sections of primary disease site and impose a transfer learning approach to predict the primary site of liver cancer *in situ*, without relying on training data drawn exclusively from liver metastases. Secondly, we observe that the first stage model may be inflexible to alternative sites of metastatic tissue. Instead of training a model to identify tiles containing cancer tissue in liver, a more generalizable model may be trained on a broad diversity of primary cancers and regularized appropriately to identify cancer independently of the host tissue. Third, although our model was shown to perform similarly to board-certified pathologists, we have not thoroughly considered the manner by which these types of deep learning models might optimally improve current workflows of practicing pathologists. We believe that robust translation of deep learning systems such as the one presented in this paper may continue to supplement and augment clinical decision-making processes dependent on medical image analysis.

## Acknowledgments

This work was supported in part by the National Cancer Institute (U54CA209988, U2CCA233280), the OHSU Center for Spatial Systems Biomedicine, the Knight Diagnostic Laboratories, and a Biomedical Innovation Program Award from the Oregon Clinical & Trantslational Research Institute. We extend our thanks to the staff at the OHSU Knight BioLibrary for their support in data access and dissemination. Further, we gratefully acknowledge the resources of the Exacloud high performance computing environment developed jointly by OHSU and Intel and the technical support of the OHSU Advanced Computing Center.

Biographies and photographs of the authors are not available.

